# Unveiling the Gut Microbiome Network Fingerprint: A Deep Learning Approach to Predict Immunotherapy Response in Melanoma Patients

**DOI:** 10.1101/2024.10.21.619471

**Authors:** Jie Sun, Zhenjiang Fan, Aditya Sriram, Hyun Jung Park

## Abstract

Immune checkpoint inhibitors (ICI) are critical in cancer therapy, harnessing the immune system to fight tumors. The gut microbiome, a diverse ecosystem of trillions of microorganisms, has emerged as a key influencer of ICI efficacy. However, the specific gut bacteria linked to ICI response in melanoma patients remain unclear, with studies showing inconsistent findings across cohorts. Attributing the lack of consensus to multiple layers of non-linear interactions with host factors and within the microbial community, we developed DeepMicroNET, a novel deep neural network (DNN) model-driven network analysis method to identify and characterize the gut bacteria associated with treatment response. Utilizing data from five cohorts of advanced melanoma patients undergoing ICI therapies, we demonstrate how DeepMicroNET identifies gut bacteria consistently predicting immunotherapy responses (response-associated or RA bacteria). Further, DeepMicroNET revealed that these bacteria are associated with key immune pathways, such as antigen processing and presentation, offering novel insights into the molecular mechanisms that could be targeted to improve therapeutic outcomes. Altogether, by identifying gut bacteria robustly associated with immunotherapy response, DeepMicroNET opens the door to a new era of personalized medicine based on the gut microbiome profile.

## Introduction

The intestinal microbiota is a mixture of microorganisms such as bacteria, viruses, archaea, and fungi. Its significance for host physiology is clear with effects on nutrient processing, drug metabolism, and immune response.^1^ Especially, due to the significant impact on the immune response, the cancer field aims to use the gut microbiome to boost immunotherapies. Immunotherapies are designed to help the immune system fight cancer using checkpoint inhibitors (ICI) such as programmed death-ligand, programmed cell death protein 1 (PD-1) and cytotoxic T-lymphocyte-associated protein (CTLA-4) monoclonal antibodies (mAbs). Recently, the gut microbiome transplantation from ICI responders have yielded durable clinical responses in nearly 40% of patients with advanced melanoma who were previously unresponsive to ICIs^2–6^. This finding links the gut microbiome to ICI therapy responses.

However, despite extensive investigation across diverse cohorts, no consensus is made on which gut bacteria are linked to ICI response in melanoma patients, hindering further translation of the findings. For instance, Studies by Matson et al. ^2^ and Gopalakrishnan et al. ^5^ reported different microbes linked to treatment response in melanoma patients. Matson et al. found a group of eight species driven by Bifidobacterium longum. Gopalakrishnan et al. found a group driven by Faecalibacterium prausnitzii. Recent meta-analyses ^7^ have further underscored these inconsistencies with strong cohort-specific effects on ICI response across melanoma patients collected from five US cities: Chicago, Dallas, Houston, New York, and Pittsburgh.

Critical examination attributes the lack of consensus to multiple levels of non-linear interactions between gut bacteria and host factors and within the microbial community. Firstly, the gut microbiome interacts with host biology nonlinearly^8^ due to several factors such as host-microbiome feedback^9^ and host genetics^10^. To identify the gut bacteria nonlinearly connected to treatment response, it is important to develop sensitive methods to capture such a nonlinear signal. For example, McCulloch et al. showed that advanced methods on batch-corrected data have improved reproducibility across cohorts. Recently, deep learning approaches have been proposed to improve performance. For example, DeepGeni, a deep autoencoder-based method^11^, predicted ICI therapy responders better than a test-based classifier^12^. Yet, autoencoders are not ideal for prediction tasks due to their unsupervised nature ^13^, spurring to explore other DNN architectures. Secondly, the gut microbiome interacts nonlinearly within the community due to the complexity of microbial ecosystems,^4^ nonlinear metabolic pathways,^5^ and mutualistic relationships^6^. Considering the interactions necessitates a shift from individual-level characterization to neighborhood-wise characterization. For instance, McCulloch et al. yielded better reproducibility across multiple cohorts not at the level of individual species, but at the level of clusters generated by k-means clustering. Previous network approaches have successfully captured microbiome interactions in diseases such as Alzheimer’s disease^2^ or type 2 diabetes (T2D)^3^. However, they rely on traditional statistical measures, which only capture linear relationships, such as correlation coefficients^23^.

To mitigate these challenges and identify the gut bacteria associated with ICI response consistently across multiple cohorts, we developed a DNN-based network analysis method, DeepMicroNET (Deep Learning-based Microbiome Network). DeepMicroNET is the first method to incorporate a DL approach for identification and network-model techniques for characterization of ICI response-associated gut bacteria. By running it on the five melanoma cohorts underlying ICIs ^2,5,14,15^ and comparing with various regression methods and DeepGeni, we will identify the ICI response-associated (RA) bacteria and validate DeepMicroNET’s DNN component. Then, we will map the interactions within the gut bacteria community to build a network model based on the interactions, with which to characterize the network property of the ICI-RA bacteria vs. other gut bacteria. This will give systematic insights into the intricate interplay of the gut bacteria shaping the ICI immunotherapy response.

## Results

### DeepMicroNET incorporates the DNN framework with network analysis to identify and characterize gut bacteria linked to treatment response

We developed DeepMicroNET using several innovative approaches. The first key innovation is in the design of supervised training where we designed a loss function to amplify the separate signal tied to treatment response or their interactions (**Fig. 1A**). This allows us to target the relevant aspects of the gut bacteria signal more effectively. This is a novel approach compared to DeepGeni, the only other deep-learning model for microbial analysis. As DeepGeni particularly relies on autoencoder to learn efficient representations of the whole aspect of the microbiome, the representation is expected to hold multifaceted signals of the microbiome, such as on nutrition processing, drug metabolism, and immune response ^1^. Thus, the autoencoder-based training may dilute the signal particular for the immune response. Second, while it is straightforward to build an unweighted network model by identifying associated bacteria pairs using a DL method and combining them, weighted network models bring greater flexibility and accuracy in modeling complex systems, providing richer insights that are difficult or impossible to obtain with unweighted models. To build weighted network models, deepMicroNET implements multiple instances of knockoff in the DNN framework. While our DNN-based causal inference method ^16^ used a single instance of knockoff to estimate the effect size and control false discovery rate (FDR), it relies on assumptions specific for causal inference (see Methods). For broader applications without the assumption, such as gut microbiome analysis, DeepMicroNET makes a novel extension with multiple knockoffs to estimate the statistical significance of the effect size (see Methods). The estimated significance helps not only prioritize the gut bacteria interactions, but also determine the weight of the bacterial interactions in a network model. In the weighted network model, deepMicroNET characterizes how effectively some gut bacteria influence the treatment responses (**Fig. 1B**) by applying various measures developed for weighted networks, such as weighted shortest path and weighted centrality (see Methods).

**Figure 1.**
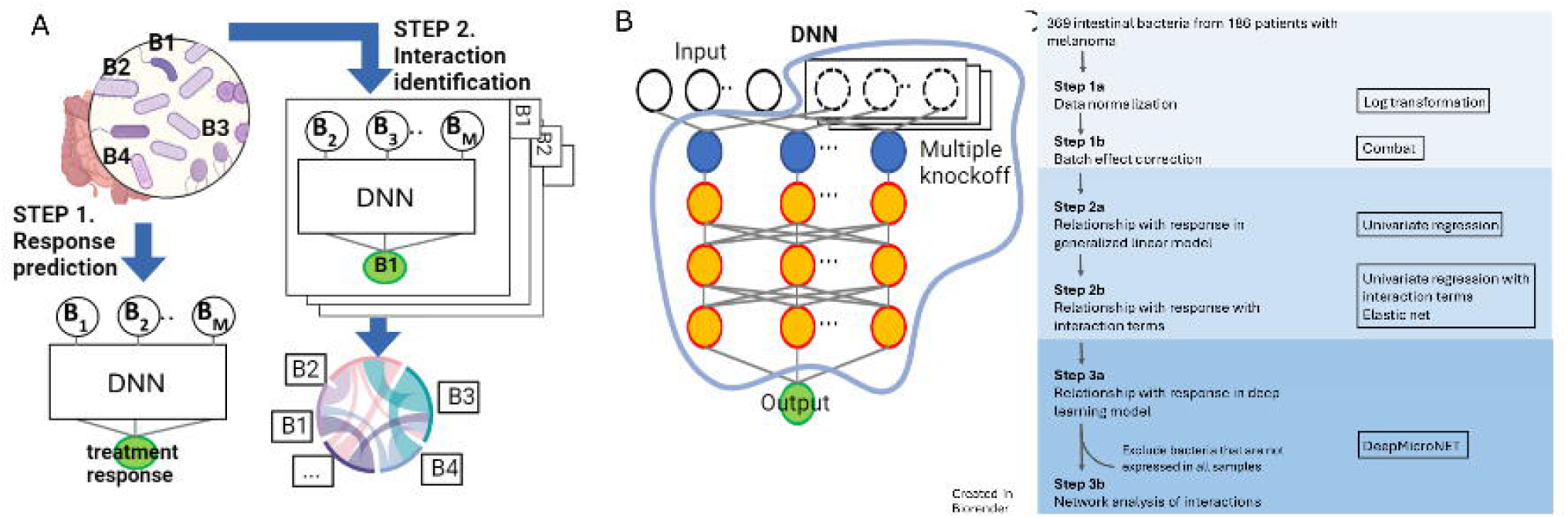
Design Principles of DeepMicroNET. **(A)** Overview of deepMicroNET algorithm that use the deep learning module differently to predict response with gut bacteria (Step 1) and to identify their interactions (Step 2). **(B)** Overview of deepMicroNET’s deep learning architecture. Multiple knockoffs are designed to estimate statistical significance of the input-output associations. **(C)** Overview of machine learning, visualization, and statistical methods applied in the present study.

### Regression analysis highlights the nonlinearity of gut bacteria interaction

To assess how linearly or nonlinearly gut bacteria interact for ICI response, we first downloaded the microbiome datasets of ICI-treated melanoma patients (**S. Fig. 1A)** from five published cohorts, and batch-corrected the data (**S. Fig. 1B, C**, see Methods) consisting of 186 sequenced samples. The definition of patients who were responders (Rs) and non-responders (NRs) was maintained as reported in the original study. Using the data, we developed multiple regression models of different complexities as follows. Firstly, we developed a logistic regression model to predict ICI therapy response in each cohort (see Methods). These regression models identified different numbers of response-associated (RA) bacteria proportional to the cohort’s size (**Fig. 2A**). For example, as the New York (n=14) and Dallas cohorts (n=14) contain the smallest cohort size, they both associated only two gut bacteria with treatment response. On the other hand, the largest cohort (n=94, Pittsburgh) identified 95 gut bacteria associated with response. Also, since most of the identified gut bacteria (127 of 134, 97.6%) are unique to a single cohort, it is possible that some of these identifications are artifacts of the sample size rather than genuine indicators of an association with therapy response. To identify and validate the RA bacteria without the sample size-related artifacts, we combined the four cohorts of smaller sizes (Dallas, Houston, Chicago, and New York, n=92) as non-Pittsburgh cohort. Then, we built a regression model using the Pittsburgh cohort (n=94) and validated the model on the non-Pittsburgh cohort. The results generally reassure the well-known lack of consistency in the RA bacteria. First, while the Pittsburgh and the non-Pitt cohorts associated 89 and 21 bacterial species with therapy response, only 7 of them are common (p-value = 0.22). Second, between the two cohorts, the significance association of the bacteria to response is slightly negatively correlated (**Fig. 2B**, Adjusted R^2^=-0.0023), indicating that the regression model yielded generally inconsistent estimates between the cohorts.

**Figure 2.**
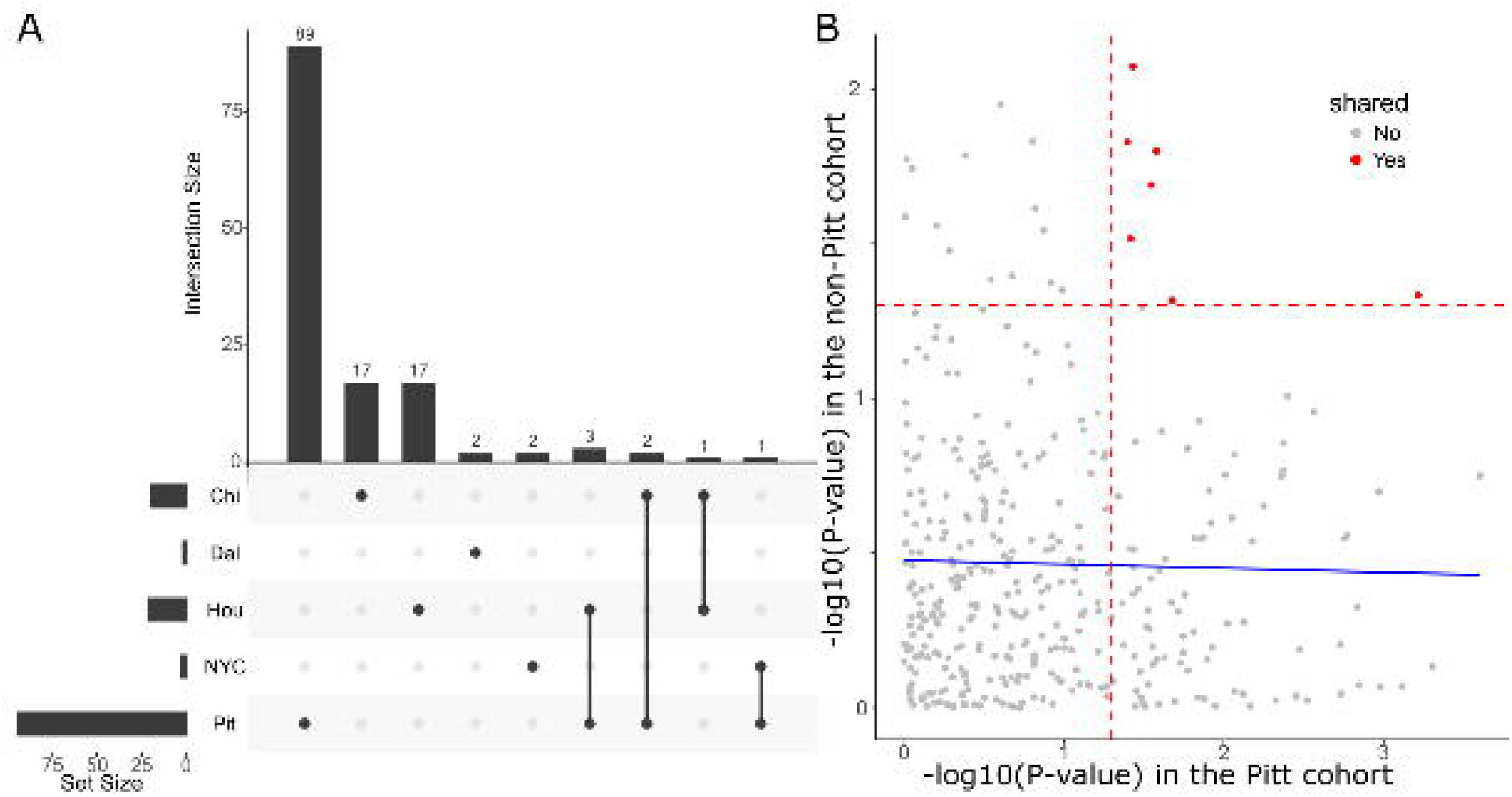
Regression analysis highlights the nonlinearity of gut bacteria interaction. **(A)** The Upset plot of the gut bacteria identified associated with response in each cohort. **(B)** Association between bacteria and response identified in the Pitt and non-Pitt cohorts. The regression line (blue line) indicates that the association between the Pitt and non-Pitt cohorts is generally uncorrelated or slightly negatively correlated. Gut bacteria significant in each cohort is in red defined by the vertical and horizontal red line in the Pitt and non-Pitt cohort.

To assess whether bacterial interactions could explain the observed inconsistency, we developed an Elastic Net model using the 7 bacteria commonly found in both the Pitt and non-Pitt cohorts, along with all pairwise interactions between them. In the Pitt cohort, the model identified 1 significant taxon and 7 significant interactions, while in the non-Pitt cohort, 4 significant taxa and 7 significant interactions were selected. However, only 1 taxon and 1 interaction were common between the two cohorts. Expectedly, when this model is trained on the Pitt cohort and tested on the non-Pitt cohort, the area under the curve (AUC) was 0.59, indicating limited predictive power. Expanding the model to include all 369 bacteria and their pairwise interactions revealed a similar trend: no overlap in the significant bacteria interactions identified between the two cohorts (17 in the Pitt cohort and 30 in the non-Pitt cohort). In another Elastic net model with all 369 bacteria with all pairwise interactions of them, the same trend was found, no overlap between 17 and 30 bacteria interactions identified in the Pitt cohort and the non-Pitt cohort, respectively. These findings suggest that the regression models struggle to capture the nonlinear dynamics of gut bacteria interactions influencing ICI responses, highlighting the need for more advanced modeling approaches.

### DeepMicroNET identifies the consistent gut bacteria effect on ICI response

To assess how consistently deepMicroNET can capture the nonlinear gut bacteria effects on the treatment response status, we first measured AUROC from the regression models we developed above, DeepGeni, and DeepMicroNET with the Pitt cohort as discovery and the non-Pitt cohort as validation cohort. As DeepGeni can utilize various machine learning (ML) models in the process, we ran it with three machine learning models, random forest (DeepGeni-RR), neural networks (DeepGeni-NN), and support vector machine (DeepGeni-SVM) (**Fig. 3A**). The results show that deepMicroNET outperforms all the other methods in AUC. The outperformance to the regression-based models indicates nonlinear relationships between the gut bacteria and the outcome. Also, the outperformance to all three deepGeni suggests that the task-specific training in DeepMicroNET adequately amplifies the signal underlying the treatment response. Second, as DeepMicroNET also estimates the association significance between each gut bacteria to response, the significance values are correlated between the two cohorts (R^2^=0.2, p-value=1.0e-19, **Fig. 3B, S. Fig. 2**). In contrast to the negative correlation from the regression model-based experiments (**Fig. 2B**), this positive correlation indicates that DeepMicroNET estimates the gut bacterial effects on ICI response consistently between the cohorts.

**Figure 3.**
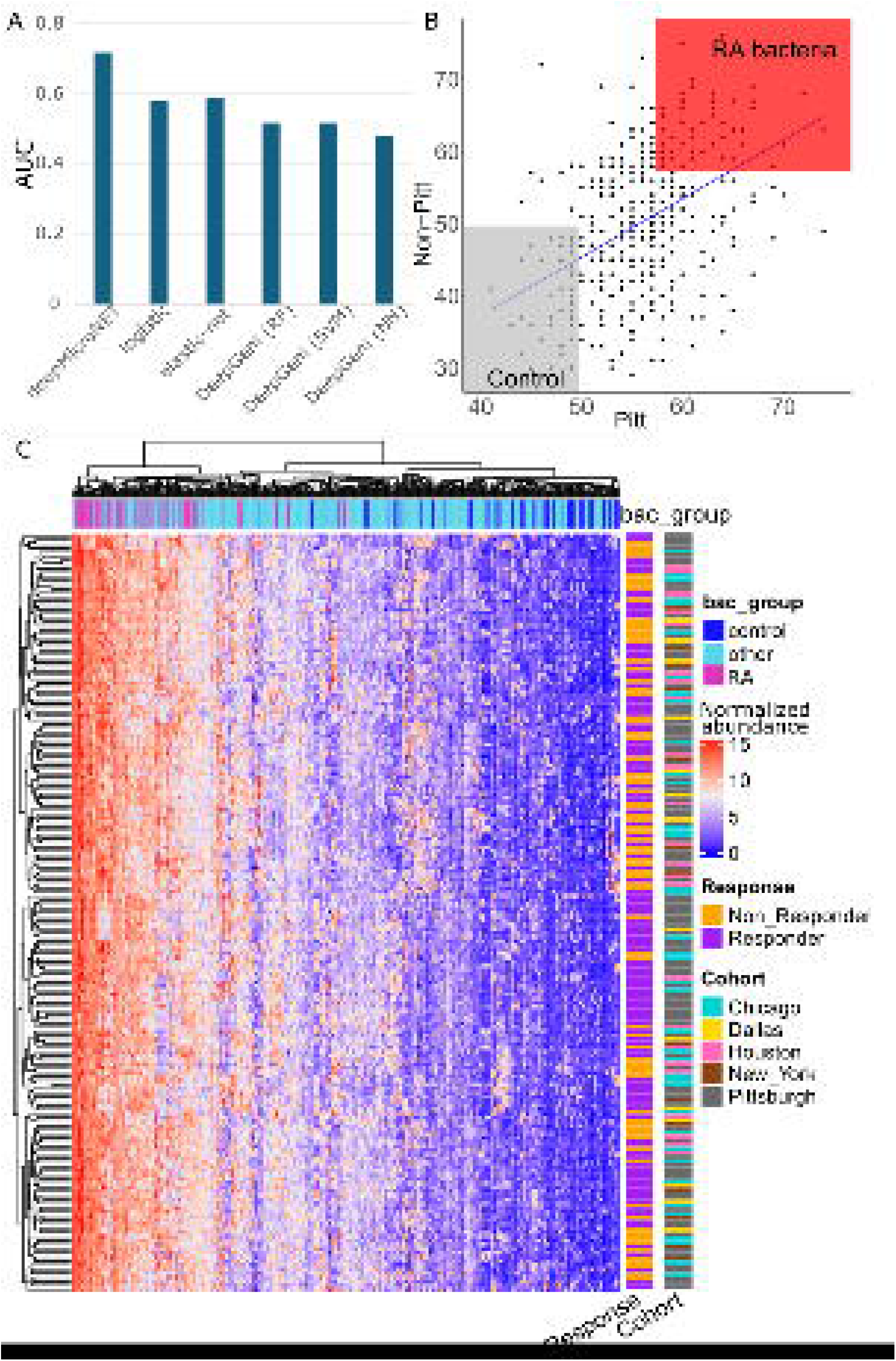
DeepMicroNET identifies the consistent gut bacteria effect on ICI response. **(A)** Comparison of deepMicroNET with regression-based (logistic and elastic-net) and deep-learning (deepGeni with random forest (RF), support vector machine (SVM), and neural networks (NN)) in predicting ICI response based on the gut microbiome data. **(B)** Significance measure of the association between each gut bacteria and treatment response identified in the Pitt (x-axis) and the non-Pitt (y-axis) cohort. 58 ICI-RA bacteria and 54 control bacteria were identified, with a similar number of gut bacteria in each group, using confidence cutoffs of 57 and 50, respectively. **(C)** Heatmap of normalized abundance values of gut microbiome of different bacteria group (bac_group: RA, control, other) sampled from different cohorts (Chicago, Dallas, Houston, New York, and Pittsburgh) of either responders or non-responders.

To further characterize the gut bacteria associated with ICI therapy response (ICI-RA bacteria), we identified them based on the significance measure estimated above. For effective characterization, we also identified those that are not associated with response based on the significance measure. Specifically, we used the significance cutoff of 0.57 and 0.5 for effective downstream analyses with similar numbers of bacteria (58 ICI-RA bacteria and 54 control bacteria). To assess if the RA bacteria are identifiable via a traditional way of looking at the gut bacteria abundance data, we ran hierarchical clustering on the batch-corrected data by bacteria and by samples (**Fig. 3C**). The clustering result shows that, while the batch effect was corrected well to get rid of cohort-specific effects, responders and non-responders do not differ in the expression profile of RA bacteria. Rather, RA bacteria are generally highly abundant in all the samples, whether responders or non-responders. Altogether, DeepMicroNET identifies the gut bacteria with strong and consistent effects on ICI therapy response, which is not straightforward if relying only on the abundance information.

### DeepMicroNET captures the host immune system affected by the gut microbiome

To examine the potential biological functions of the RA bacteria, we further performed taxon set enrichment analysis (TSEA) on the RA vs. the control bacteria. TSEA is a module implemented in MicrobiomeAnalyzer^18^ that characterizes taxonomic signatures by linking microbial taxa to genetics attributes. The TSEA result showed that the RA and the control bacteria groups are enriched with different gene sets with no intersection between them (**S. Fig. 3A, S. Fig. 3B**). The genes uniquely enriched with ICI-RA bacteria have been associated with better responses to ICI therapies. For instance, the HLA-mediated immunity, involving HLA-DOA gene, is linked to ICI response in pan-cancer patients^19^. Specifically, for melanoma, immune-related genes such as IL34^20^ and ERAP2^21^, which are involved in modulating immune responses, have also been associated with more favorable outcomes in immunotherapy.

To further examine the functional differences between the bacteria groups, we performed the Ingenuity Pathway Analysis (IPA) comparing the gene sets enriched for the two bacteria groups. First, the IPA comparison analysis under Diseases & Functions terms clearly showed that the bacteria groups show the biggest enrichment difference in melanoma, among more than 200 cancer terms (**Fig. 4A, S. Table 1**). This result suggests that the RA bacteria may have a unique impact on melanoma progression, rather than on cancer in general. Second, based on the IPA comparison under canonical pathway terms, antigen presentation and class I MHC mediated antigen processing terms were found among the top four most enrichment p value difference between the groups (**Fig. 4B, S. Table 2**). Antigen presentation and processing are critical components for the human immune system, such as cytotoxic T cells and regulatory T cells, to fight cancer^1^ and thus have been positively correlated with better responses to ICIs.^22^ Thus, the results suggest that the ICI RA gut bacteria are causally associated with treatment response for melanoma patients through modulating the immune system.

**Figure 4.**
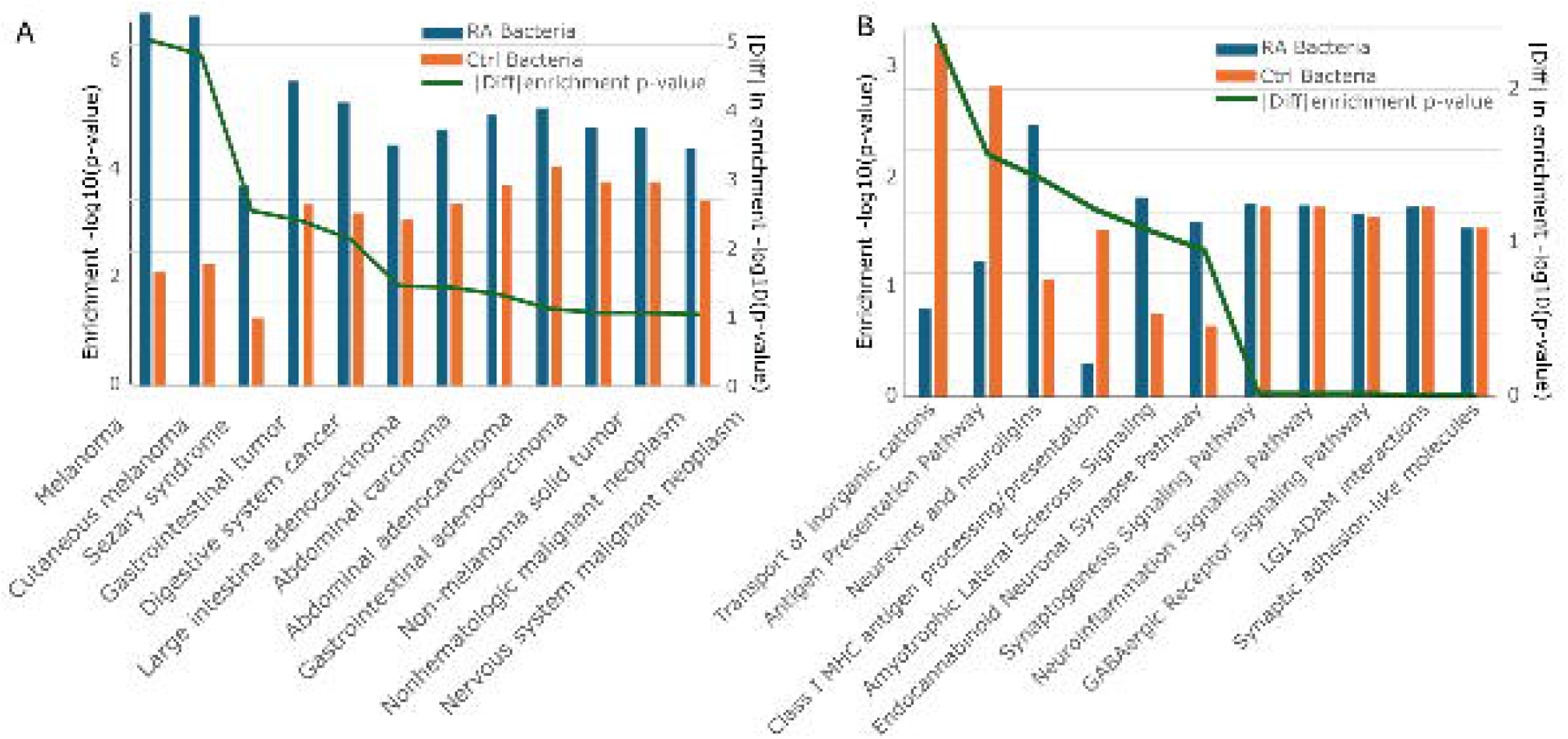
DeepMicroNET captures the host immune system affected by the gut microbiome. **(A)** Ingenuity Pathway Analysis (IPA) under the “Diseases & Functions” category for the RA and the control bacteria. The blue bars and the orange bars represent the enrichment significance of the RA and the control bacteria group. The green line represents the absolute value of the difference in the enrichment significance for the RA and the control bacteria. The terms were ranked by the absolute enrichment significance difference. **(B)** IPA canonical pathway analysis for the RA and the control bacteria. The blue bars and the orange bars represent the enrichment significance of the RA and the control bacteria group. The green line represents the absolute value of the difference in the enrichment significance for the RNA and the control bacteria. The terms were ranked by the absolute enrichment significance difference.

### Network analysis captures the complexity of gut bacteria interactions

To elucidate how the RA bacteria group efficiently affects the immune system and elicit responses, we combined the Pittsburgh and non-Pittsburgh cohorts and selected 138 abundant bacteria out of 369 bacteria (52 RA and 86 non-RA bacteria). On the data, we ran DeepMicroNET to identify 1,578 nonlinear relationships based on the effect size (see Methods, **Fig. 5A, S. Table 3)**. Close examination of the network model allows us to examine how RA bacteria interact for ICI response. First, the RA bacteria are generally more highly connected than the non-RA bacteria (**Fig. 5B**). Consistently, RA bacteria communicate with other bacteria generally in shorter path distances (**Fig. 5C**). Since the shortest path distance represents the most efficient route for signals to propagate, RA bacteria are expected to affect host biology through efficient communications among the gut bacteria. Other metrics, such as betweenness centrality and closeness centrality, reveal the same characteristics about the RA bacteria being highly connected and efficiently influencing the microbiome (**S. Fig. 4**). Second, the RA bacteria are not different than the other bacteria in terms of clustering coefficients (**Fig. 5D**). Since clustering coefficient is the fraction of pairs of each node’s neighbors that are connected to each other, 0 means none of the neighbors are connected and 1 means all neighbors are connected for a particular node. Since the values indicate varying levels of network cohesion or community structure between groups in a network^23^, a similar distribution of clustering coefficients between the RA and the other bacteria group suggests that they are spread or concentrated equally across multiple gut bacteria communities. We further compared the RA and non-RA bacteria group by the relationship between clustering coefficients and degree (**Fig. 5E**). While the non-RA bacteria group does not show a clear pattern, the clustering coefficient is significantly (p-value=0.003) negatively correlated with degree in the RA group. This negative correlation indicates that hub bacteria among the RA group connect diverse groups of bacteria that may not interact with each other, while low-degree bacteria form tightly-knit functional clusters. This structure supports the modularity and functional specialization of the microbiome, with the hub RA bacteria serving as generalists linking different groups and the other RA bacteria specializing in local cooperative functions. Altogether, the gut bacteria associated with treatment response exert stronger effect on the microbiome but equally spread across, resulting in efficient influence on the gut microbiome community.

**Figure 5.**
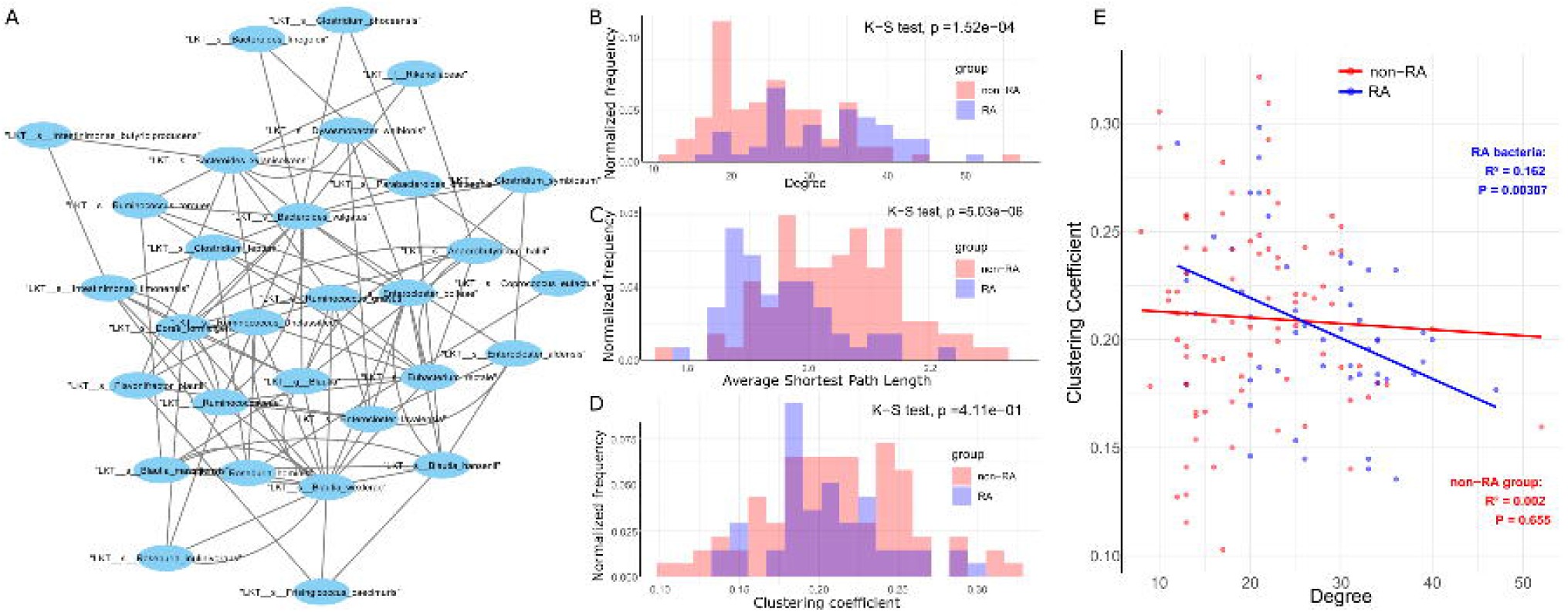
Network analysis captures the complexity of gut bacteria interactions. **(A)** A network was constructed using 52 RA bacteria and 86 non-RA bacteria, with the effect size calculated by DeepMicroNET as interaction weights. **(B)** The degree distribution of each node within the RA and non-RA bacteria groups. The normalized frequency was calculated by using the frequency of each bin divided by the total frequency in each group. The same technique was applied to other histograms for network property visualization. **(C)** The average shortest path length distribution of each node within the RA and non-RA bacteria groups. **(D)** The clustering coefficient distribution of each node within the RA and non-RA bacteria groups. (E) Scatter plot of the clustering coefficient of bacteria nodes in the network against the degree. The RA and the non-Ra bacteria groups are in blue and red, respectively. Similarly, the blue and red lines indicate their linearly regressed trend of the bacteria group.

## Discussion

Analyzing the gut microbiome has become crucial in understanding the efficacy of tumor immune checkpoint inhibitor (ICI) therapies. In this manuscript, we showed that traditional statistical models, while useful, struggle to capture the complex and nonlinear relationships between gut microbiota and therapeutic outcomes. In contrast, our method, DeepMicroNET, excels in identifying ICI therapy response-associated gut bacteria by developing a novel DNN module and constructing a weighted network of microbial interactions. The DNN module in DeepMicroNET improves the prediction performance of the previous DNN method, DeepGeni, by employing a task-specific loss function. DeepGeni is designed to represent the whole aspect of the data in a lower-dimensional space. However, since the whole aspect of the gut microbiome should reflect the multifaceted aspects of human life, not only a trait of our interest, our analysis reveals that amplifying the trait-specific signal, using a tailored loss function, is key to improving performance. To construct a weighted network model, DeepMicroNET innovatively designs the multiple knockoff framework and estimates the statistical significance of the microbial interactions. This innovative approach provides a detailed and systematic understanding of how microbial interactions influence therapeutic outcomes.

DeepMicroNET not only deepens our understanding of the microbial effects on the ICI therapeutic outcomes but also reveals novel insights into the molecular mechanisms that could be targeted to improve treatments. For example, the ICI therapy response-associated (RA) and the control bacteria group show the biggest difference in enriching for antigen presentation and class I MHC mediated antigen processing (**Fig. 4B**). The antigen presentation process stimulates immature T cells to differentiate into critical components of the adaptive immune system, either cytotoxic T cells (CD8+ T cells) or T helper (Th) cells (CD4+ T cells).^24^ While the original study linked cytotoxic and regulatory T cells to immunotherapy responses using single-cell RNA-Seq data,^25^ deepMicroNET allows us to find this implication only using the microbiome data. This result demonstrates the potential value of DeepMicroNET in understanding microbiome dynamics without additional host data.

DeepMicroNET further offers clinical implications. For example, the shorter path distances of ICI-RA bacteria in the microbiome implies how effectively the perturbations (e.g., modulating their abundance) on the ICI-RA bacteria might propagate through the network, suggesting that the ICI-RA bacteria can be efficient drug targets. Furthermore, the negative relationship between clustering coefficient and degree characterizes the scale-free structure of the gut microbiome relationship. This scale-free structure can contribute to the stability and resilience of the gut microbiome, as hub bacteria might help maintain functional connections between different microbial communities, even if individual bacteria within communities are lost or disturbed.

Despite the clear implications, the current study has some limitations. We used the total of 168 melanoma patients collected from five US cities without cohorts from similar international trials (e.g. UK and Netherlands^26^). This is to minimize the international level of dietary and environmental confounders, which are high in gut microbiome data and could bias the analysis.^27^ However, the cohort data presents some inherent challenges in sequencing technique difference and the imbalance of responders and non-responders. First, the data we used were generated by two different techniques, 16S rRNA and Shotgun metagenomic sequencing. Despite the technical difference, a previous meta-analysis found similar results between these 16S and shotgun sequencing data ^2^. However, further study is needed to comprehensively understand the impact of the difference. Additionally, some cohorts, such as the Pitt cohort, have more responders (68) than non-responders (26), which raises concerns about overfitting to the response mechanism. In the future, we plan to resolve these issues by collecting more melanoma cohorts and carefully correcting the batch effects with respect to the limitations.

Methodological limitations lie in improving its statistical significance measures and expanding network metrics. While the comparison of response-associated (RA) and control bacteria offers insights into the gut bacteria dynamics, the lack of a defined empirical distribution for the significance measure remains an issue to identify significant RA bacteria. This is due to the current implementation training the model serially, each with a single instance of knockoff. In the future, we plan to introduce parallel processing with multiple knockoffs to efficiently establish the empirical distribution. Also, to better understand microbial interaction patterns, we will implement other weighted network measures, e.g., network motifs and rich club coefficients. Network motifs are small, recurring subgraphs or patterns within a larger network. Identifying motifs in the microbial interaction network will help reveal structural features that may indicate key functional roles in the microbial community. The rich club coefficient measures whether high-degree nodes in a network tend to form a tightly interconnected “club.” Since the ICI RA bacteria are high-degree nodes in the network, this metric will examine the level of connectivity among the RA bacteria.

Our findings have real-world potential that could transform cancer immunotherapy. By identifying gut bacteria linked to immunotherapy response, we pave the way for personalized medicine based on the gut microbiome. In the near future, microbiome profiling could become a standard part of cancer treatment, helping clinicians predict which patients will respond to immunotherapy. This approach could result in more targeted treatments, sparing non-responders from unnecessary side effects and lowering healthcare costs.

## Methods

### Data collection and batch effect removal

Data was collected from five published datasets of different sample size across five locations in the US: Dallas (n=14), Chicago (n=39), Houston (n=25), New York (n=14), and Pittsburgh (n=94). To make the dataset comparable in a single study, we normalized and performed the batch-effect-correction on the data sets according to the pipeline developed for this data type.^29^ The data was normalized using log2 and quantile transformations. To address batch effects between datasets, we applied the Combat R package (v. 3.20.0) ^30^ and used UMAP to visualize differences before and after batch correction.

To ensure comparisons with enough sample size, we combined datasets from Dallas, Chicago, Houston, and New York into a non-Pitt cohort (n=92), which included 43 responders and 49 non-responders. The Pittsburgh cohort contained 68 responders and 26 non-responders. In total, 369 bacterial species were analyzed for each patient.

### Regression models and elastic nets

To investigate whether the relationship between microbiome composition and ICI response in melanoma patients is linear, we first established a univariate regression model, using response as the dependent variable and each bacterium as an independent variable. After identifying seven statistically significant bacteria (FDR < 0.05) shared between the Pitt and non-Pitt cohorts, we applied a hypergeometric test to assess whether this overlap occurred by chance.

We further explored the potential linear relationship by using elastic net models. Interaction terms were added to the elastic net models, incorporating the seven bacteria commonly found in both the Pittsburgh and non-Pittsburgh cohorts, along with all pairwise interactions among them. The models were trained using the glmnet package in R^31^.

### DeepMicroNET model development: multiple knockoff

We developed DeepMicroNET by extending from DeepVASE^16^ to estimate prediction confidence. In our earlier DNN model, predictions exhibited some randomness due to the use of “glorot_normal” as the Kernel Initializer, which initializes the neural network weights. To obtain consistent results, we previously set the seed in the experiments. Previous studies proved that this knockoff filter controls a slightly modified version of FDR under mild conditions for causal inference^3^. Specifically, the mild conditions are about the joint distribution of input data where the Markov blanket is well defined within the data and unique so that we have a cleanly stated selection problem ^4^. However, the assumption of Markov blanket posits that the features are connected only to exactly represent their conditional independence relationship. The concept of a Markov blanket is central to causal inference because it helps isolate the direct causes, effects, and indirect paths through which information about the variable of interest flows. However, it is not known if this assumption holds for gut microbiome data analysis. To estimate statistical significance without the Markov blanket assumption, we improved the DNN model by using multiple (in this study, 100) different seed settings, counting how often features were selected across 100 iterations, the counts of which was defined as “prediction confidence”. This parameter can be adjusted for 1000, 2000, or more iterations as needed.

### DeepMicroNET model development: deep-learning framework

Our DNN architecture consists of an input layer with twice the number of neurons as input features, accommodating both the original features and knockoff features in a pairwise manner. The network has two hidden layers, each with the same number of neurons as the input variables and uses the rectified linear unit (ReLU) activation function. The weights in the network are initialized using the Glorot normal initializer, and L1 (LASSO) regularization is applied to prevent overfitting. The model is optimized using the Adam algorithm with a learning rate of 0.001, and the mean squared error (MSE) serves as the loss function during training.

To identify nonlinear associations between input variables *x*_*i*_, DAG-deepVASE constructs a series of perceptron layers between *X*_\*i*_ *=* {*x*_*1*_, *x*_*2*_,…, *x*_*i*-1_, *x*_*i*+1_,*…, x*_*M*_} and *x*_*i*_. The aim is to estimate the effect size of the association between *x*_*j*_ ∈ *X*_\*i*_ and *x*_*i*_ using model-X knockoffs. For the input variables *x*_*i*_ and *x*_*j*_, the method ensures the exchangeability property 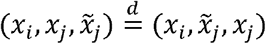, where 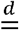 denotes equality in distribution. This property allows the identification of causal relationships, differentiating them from simple correlations.

For instance, if *(x*_*i*_, *x*_*k*_) is a correlation without a causal link, then the feature exchangeability 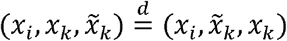 will hold, making the relationship measures |*R*_*ik*_| and 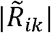 symmetric and exchangeable around zero where *R*_*ik*_ is the correlation between the original feature *X*_*i*_ and the outcome after training, and 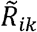 is the correlation between the knockoff 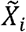 and the outcome after training. On the other hand, if (*x*_*i*_,*x*_*j*_) is a causal relationship, 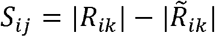 will capture the deviation of this relationship from the null hypothesis. The knockoff matrix 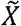 is designed to match the correlation structure within *X* while minimizing cross-correlation with the outcome variable *Y*.

The knockoff variables 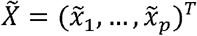 follow two main properties: (i) Exchangeability, where 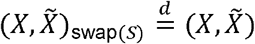, where swapping denotes exchanging any subset *S* ⊆ {1, …, *M*} of the variables *x*_*j*_, and (ii) Independence, meaning 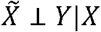, ensuring that 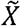 is independent of *X* given the outcome *Y*.

To construct knockoffs, we assume *X* ∼ *N*(0, Σ), with Σ ∈ ℝ ^*M*×*M*^ as the covariance matrix. A valid knockoff construction for 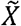 is given by:

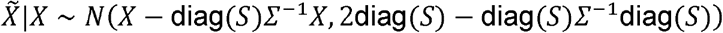

The effect size *S*_*ik*_ is computed as the difference in correlation strength between the original feature and the knockoff feature:

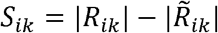

If the original feature has a stronger correlation than its knockoff counterpart, it is called as a selected feature for continued analysis. Effect size calculations for each identified association involved pairing each input variable with its knockoff counterpart and using matrix operations across the DNN’s layers to quantify the importance of each original variable relative to its knockoff. The effect size determining process utilizes weight matrices to compute feature importance scores. These importance scores are then used to calculate an effect size for each feature that reflects the strength of each feature’s association with the outcome of interest, calculated as the difference between the importance of the original feature and its knockoff variable. We parameterized *q* as a user-defined nominal false discovery rate (FDR), and subsequently set FDR to a level *q* = 0.05 to enable balance estimation of associations.

### Identifying and characterizing ICI-RA and control bacteria groups

To identify bacteria consistently selected across the Pitt and non-Pitt cohorts, we focused on a similar number of gut bacteria associated with the response (58 ICI-RA bacteria) and those not associated with the response (54 control bacteria), using confidence cutoffs of 57 and 50 out of 100 iterations, respectively.

To further investigate the biological functions of the RA and control bacteria groups, we utilized taxon set enrichment analysis (TSEA) within MicrobiomeAnalyst^18^. Following this, we conducted functional enrichment analysis on the gene sets enriched in these two bacteria groups using Ingenuity Pathway Analysis (IPA), with a particular focus on Diseases & Functions and IPA canonical pathway terms.

To evaluate the predictive performance of DeepMicroNET against other models, including logistic regression, elastic net, and DeepGeni, we used the Pitt cohort as the training set and the non-Pitt cohort as the test set. We calculated AUROC values for each model to assess their accuracy in predicting outcomes. The logistic regression and elastic net models used clinical response as the outcome variable, with the seven significant taxa identified in both Pitt and non-Pitt cohorts through univariate regression, along with their pairwise interactions, as the independent variables.

### Network analysis

To maximize the sample size for this analysis, the Pitt and non-Pitt cohorts were combined. To explore the gut microbiome network, we selected 138 taxa out of 369 that were expressed across all samples. DeepMicroNET was then applied, identifying 1,578 significant (FDR < 0.01 and w score > 0.0001) nonlinear interactions among the 138 selected taxa. The resulting network model included 52 RA bacteria and 81 non-RA bacteria.

We used Cytoscape^32^ to visualize the network and calculate various network properties, including degree, average shortest path length, betweenness centrality, closeness centrality, and clustering coefficient. To enhance readability, we included only the nodes selected by the MCODE tool, which identifies highly interconnected regions within the network. The width of each edge represents the interaction weights generated by the DNN component of DeepMicroNET. The Kolmogorov-Smirnov (K-S) test was employed to compare these network properties between the RA and non-RA bacteria groups.

## Figure Captions

**Supplementary Figure 1.**
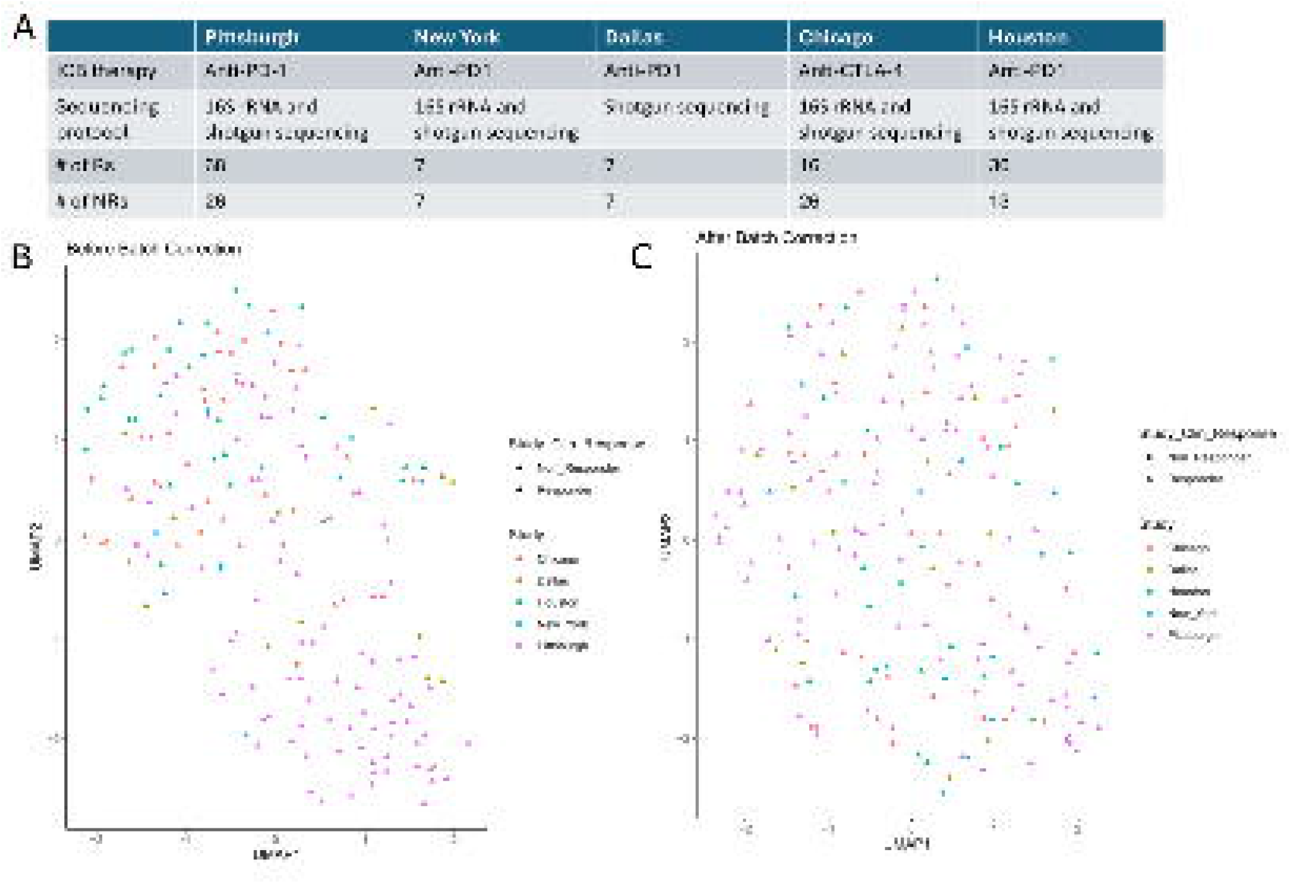
**(A)** Meta data table of the five cohorts. **(B)** Before and **(C)** after the batch effect correction across the five cohort data in the UMAP visualizations.

**Supplementary Figure 2.**
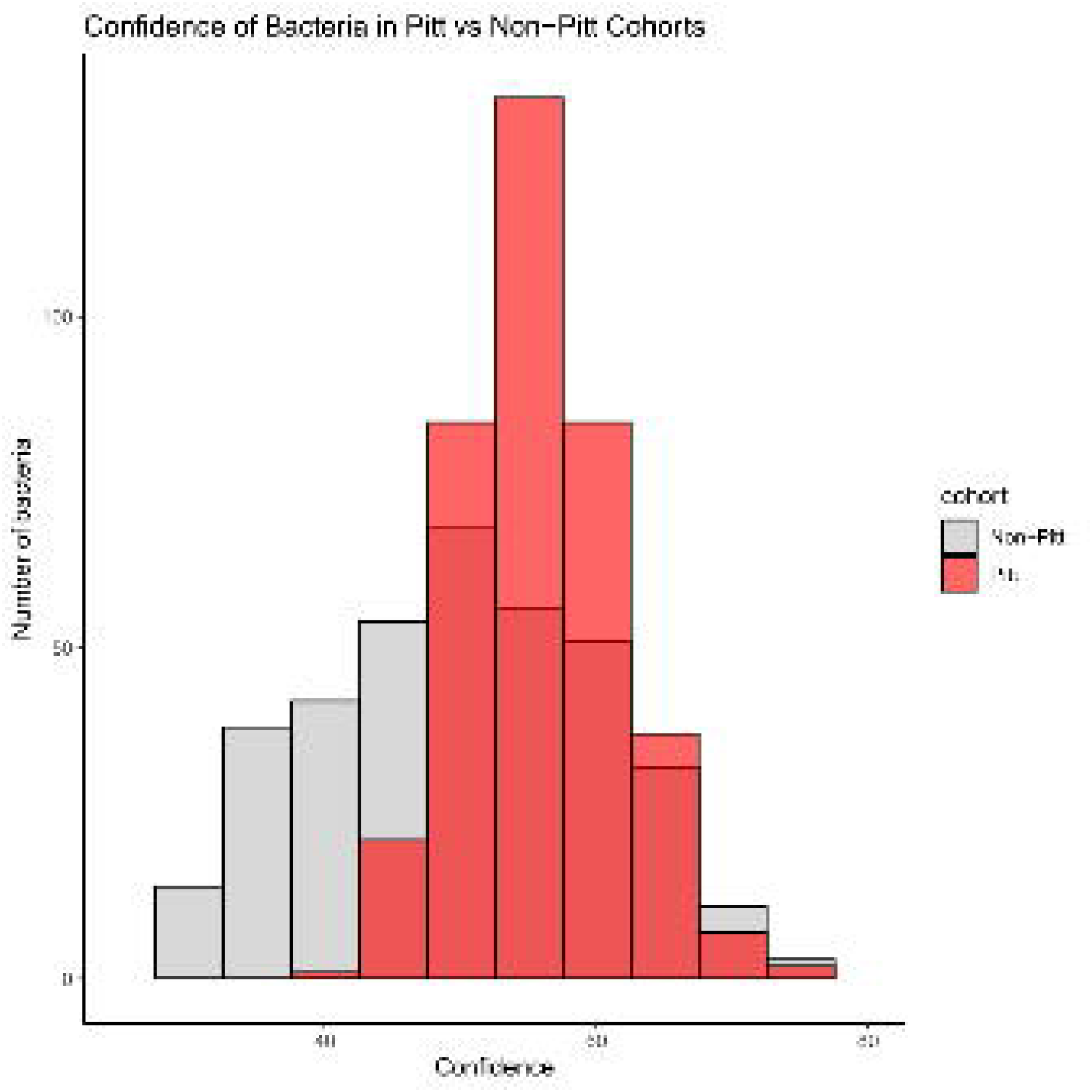
Results from DNN Component of DeepMicroNET. The x-axis of the histogram represents the association confidence level, indicating how many times the same bacteria were selected during the 100 repetitions. The homogeneous Pitt cohort has generally higher prediction confidence.

**Supplementary Figure 3.**
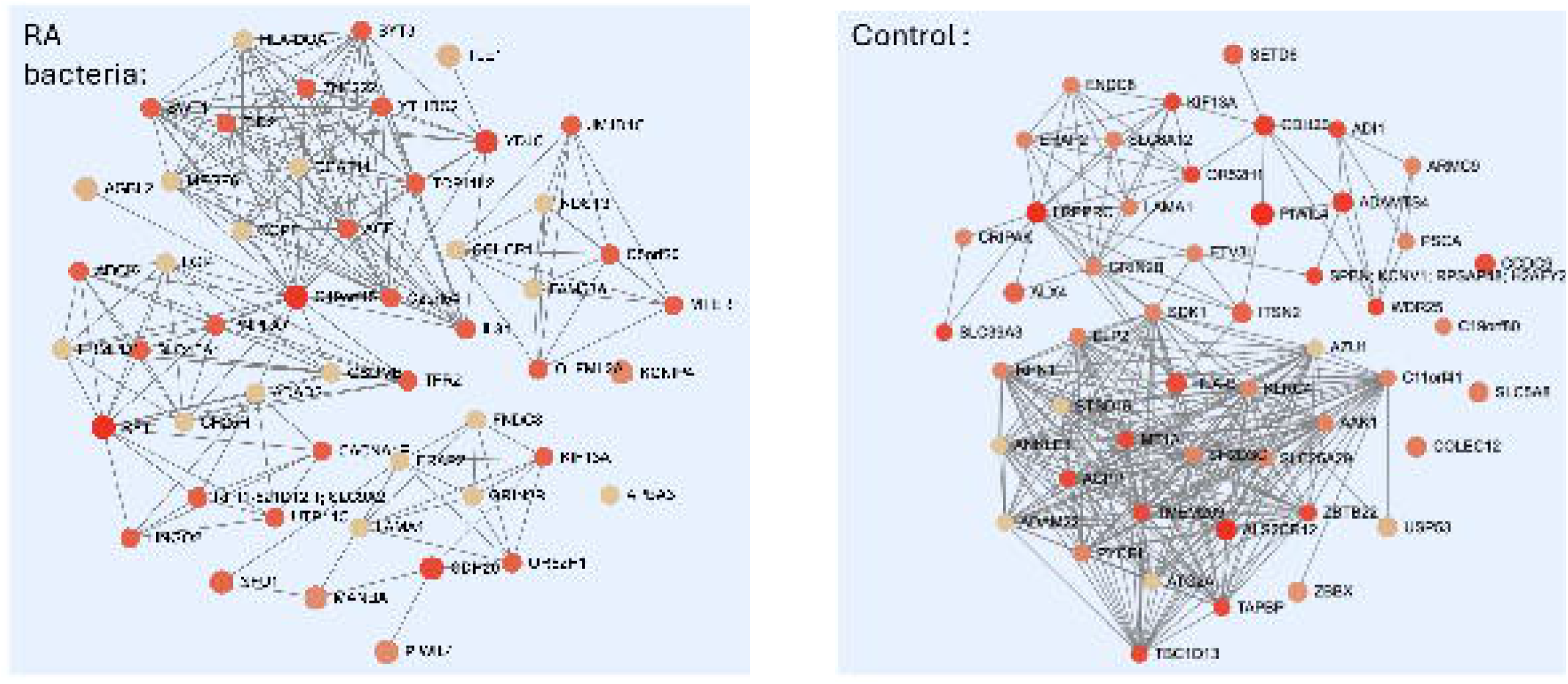
The TSEA analysis results of ICI-RA bacteria group **(A)** and control group **(B)**.

**Supplementary Figure 4.**
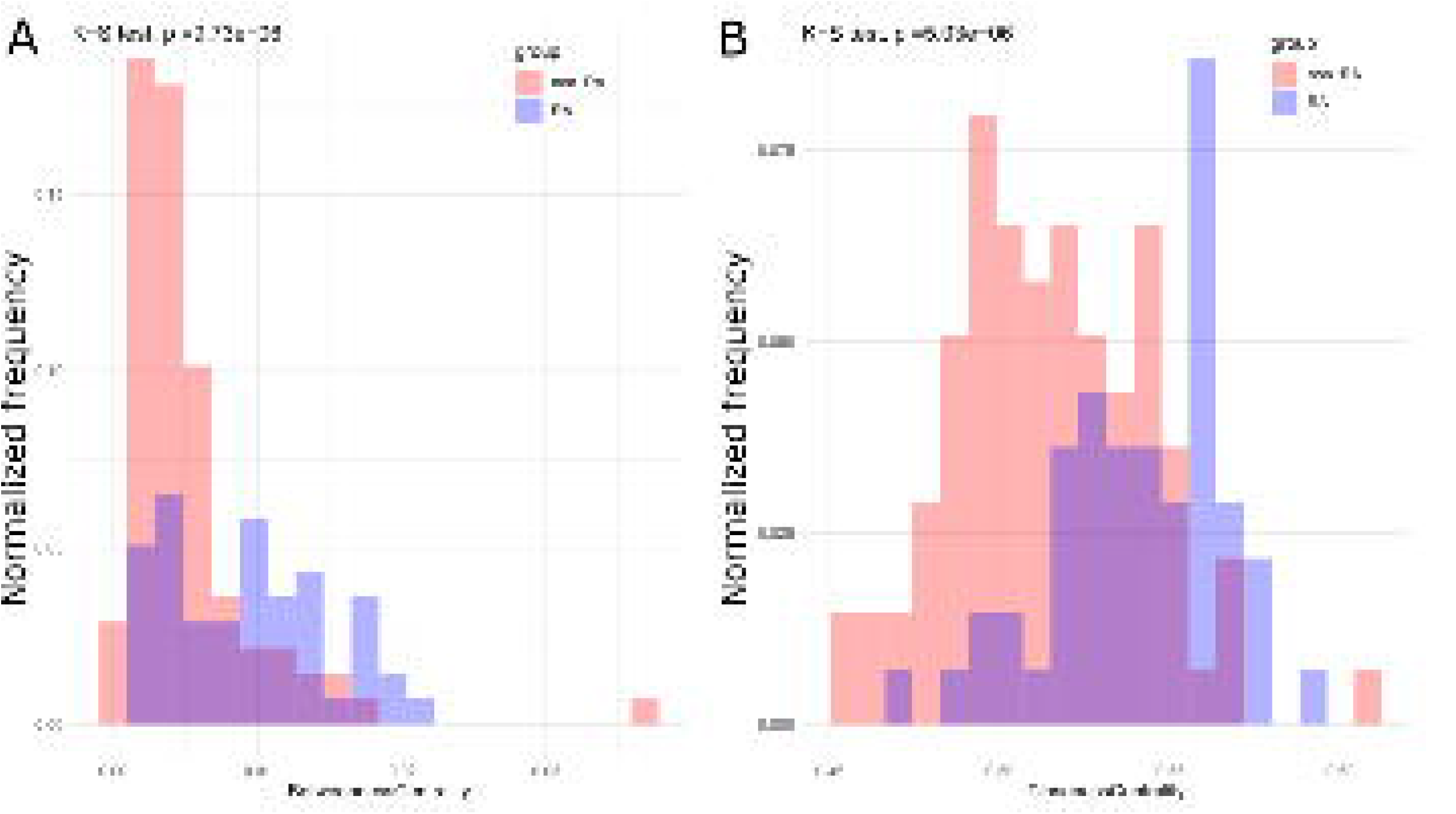
The betweenness centrality **(A)** and closeness centrality **(B)** distribution of each node within the ICI-RA and non-RA bacteria groups.

**Supplementary Table 1**. IPA diseases and functions analysis for the RA and the control bacteria.

**Supplementary Table 2**. IPA canonical pathways analysis for the RA and the control bacteria.

**Supplementary Table 3**. Various network metrics estimated for the RA and the non-RA bacteria group.

